# CATEomics: model selection and ensembling for heterogeneous treatment effect estimation from high-dimensional omics data

**DOI:** 10.64898/2026.11.16.767168

**Authors:** Takeru Furuta, Koichi Himori, Bingyuan Zhang, Akihiro Kitani, Yusuke Matsui

## Abstract

**Summary:** `CATEomics` addresses the lack of a standardized and reproducible process for choosing among conditional average treatment effect (CATE) estimators in high-dimensional omics studies, where their estimation errors cannot be evaluated directly from observed outcomes. The R package accepts a `SummarizedExperiment`, applies common preprocessing and data partitions across R and Python estimators, compares their out-of-fold estimates using proxy losses, supports model selection and ensembling, and returns the resulting CATE estimates to the original object for downstream omics analysis.

**Availability and implementation:** `CATEomics` is implemented in R and is freely available under the MIT license at https://github.com/matsui-lab/CATEomics. The software version described here (v0.99.0) is archived on Zenodo at https://doi.org/10.5281/zenodo.24189754. A vignette illustrates the complete workflow from a `SummarizedExperiment` to downstream analysis.

**Supplementary information:** Supplementary material is provided with this preprint.

## 1 Introduction

High-throughput omics assays measure thousands of molecular features in samples from individual patients. Although such data are widely used to predict prognosis, treatment selection requires knowledge of the treatment effect, that is, the expected difference in outcome between treatments for patients with a given molecular profile. Under the assumptions required for causal identification, this difference is known as the conditional average treatment effect (CATE). Estimating CATE from omics data may identify molecular profiles associated with differential treatment benefit (Zhang et al. 2017).

Many CATE estimators have been proposed (Zhang et al. 2022), but their relative performance depends on how well the assumptions of each estimator match the structure of the data, such as the outcome structure and the strength and form of treatment-effect heterogeneity (Curth and van der Schaar 2023; Mahajan et al. 2024). This dependence is especially problematic in high-dimensional omics studies, where molecular features often outnumber samples and estimates therefore depend strongly on each estimator’s modeling assumptions. Moreover, because each patient reveals an outcome under only one treatment, the true CATE estimation error is unobservable (Fig. 1A). This further complicates model assessment. Instead, candidate models can be compared using proxy losses such as the R-loss, calculated on patients not used for model fitting, to guide model selection or ensembling (Nie and Wager 2021; Mahajan et al. 2024). A valid and reproducible comparison therefore requires common preprocessing, data partitions and scoring procedures across candidate models.

**Fig. 1.**
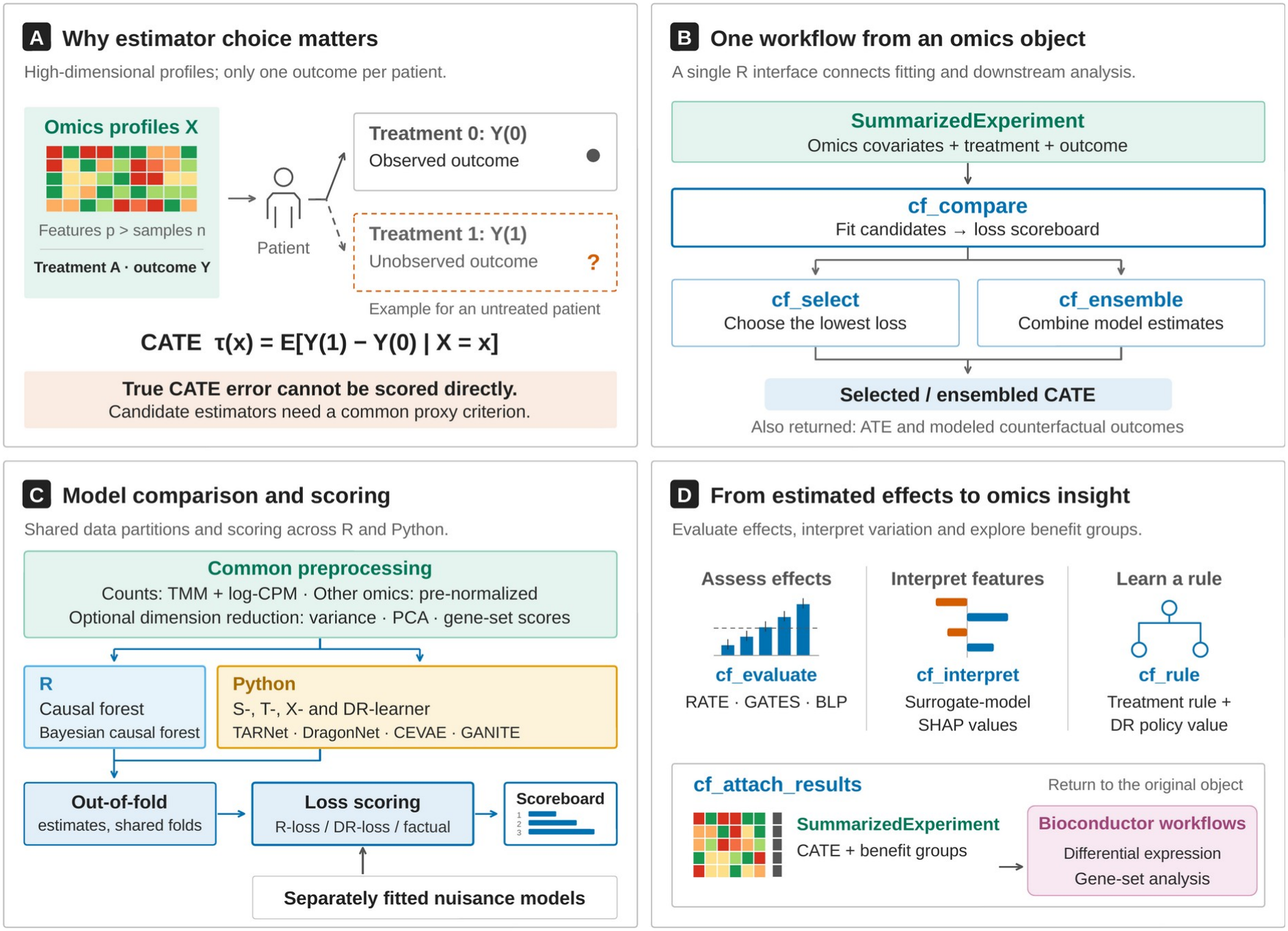
Overview of the `CATEomics` workflow. (A) Omics studies provide high-dimensional molecular features, treatment assignments and observed outcomes. Only one potential outcome is observed for each patient, which prevents direct evaluation of the true conditional average treatment effect (CATE) estimation error. (B) Starting from a `SummarizedExperiment, cf_compare` fits the candidate models and returns their losses as a scoreboard; `cf_select` chooses the model with the lowest loss, whereas `cf_ensemble` combines model estimates. (C) Model comparison and scoring. R and Python estimators share the same preprocessing (including optional dimension reduction) and data partitions. Out-of-fold CATE estimates are scored using the R-loss, DR-loss and factual loss; the nuisance models for these losses are fitted separately from the candidate models. (D) The selected or ensembled estimates are evaluated against the pseudo-outcome (RATE, GATES and BLP), interpreted with surrogate-model SHAP values and used to construct treatment rules. `cf_attach_results` writes the CATE estimates and the benefit groups defined by `cf_benefit_groups` back to the original `SummarizedExperiment` for downstream Bioconductor workflows, including differential expression and gene-set analysis. Plots and icons are schematic and do not represent empirical results.

Available software covers only parts of this workflow. `causalToolbox` provides an R interface to multiple CATE estimators, whereas `EconML` supports CATE estimation, R-loss-based model selection and ensembling in Python (Künzel et al. 2019a; Battocchi et al. 2019). Neither provides an end-to-end Bioconductor workflow that maintains consistent preprocessing and evaluation and returns CATE estimates to standard omics data containers for direct use in established workflows, such as differential expression and gene-set analysis. Estimator comparison therefore remains dependent on custom cross-language code and is difficult to reproduce.

We developed `CATEomics` to provide this workflow (Fig. 1). Starting from a `SummarizedExperiment`, `CATEomics` applies common preprocessing and data partitions to candidate R and Python estimators, estimates the effect for each patient with models fitted without that patient (out-of-fold estimates), compares these estimates using proxy losses based on auxiliary models (nuisance models) fitted separately from the candidates, and returns selected or ensembled CATE estimates and benefit groups to the original object.

## 2 Implementation

### 2.1 Overview

`CATEomics` takes a `SummarizedExperiment`, a standard Bioconductor data container (Huber et al. 2015), as its primary input and also accepts a `data.frame` or a `MultiAssayExperiment` (Ramos et al. 2017). Users specify the outcome, the treatment, the clinical covariates and the omics features, all of which are taken from a single object. Continuous and binary outcomes are supported, with effects on binary outcomes estimated as risk differences. `cf_compare` fits the candidate models and returns their losses as a scoreboard. Using this scoreboard, `cf_select` chooses a single model or `cf_ensemble` combines several (Fig. 1B; Section 2.2). In either case, `CATEomics` returns the per-patient CATE estimates 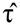, the average treatment effect and the modeled counterfactual outcomes. The CATE estimates are then evaluated, interpreted and used for downstream analysis, and the estimates and benefit groups can be written back to the original object (Fig. 1D; Section 2.3). The backend, random seed and software versions of each model are recorded for reproducibility.

### 2.2 Model comparison, selection and ensembling

Users first choose the candidate models from ten methods that `cf_compare` accepts through a single interface: R implementations of the causal forest (Athey et al. 2019) and the Bayesian causal forest (Hahn et al. 2020), and Python implementations of the S-, T- and X-learners (Künzel et al. 2019b), the DR-learner (Kennedy 2023) and four deep-learning methods, namely TARNet (Shalit et al. 2017), DragonNet (Shi et al. 2019), CEVAE (Louizos et al. 2017) and GANITE (Yoon et al. 2018), using the implementations in `EconML` (Battocchi et al. 2019), `CATENets` (Curth and van der Schaar 2021) and `Pyro` (Bingham et al. 2019) where available. The Python environment is configured automatically. For the S-, T-, X- and DR-learners, users can also choose the base learners used within each method from linear and logistic regression, penalized regression, random forests and gradient boosting.

`cf_compare` applies common preprocessing to all candidate models (Fig. 1C). Raw RNA-seq counts can be TMM-normalized (Robinson and Oshlack 2010) and log-CPM transformed internally using `edgeR` (Chen et al. 2025), whereas other omics data should be normalized before input. Users can reduce the number of features by selecting the most variable ones, by principal component analysis or, for gene expression data, by gene-set scoring (Hänzelmann et al. 2013); dimension reduction can also be omitted.

`cf_compare` then obtains out-of-fold estimates for each model, using data partitions shared by all candidate models. Using nuisance models fitted separately from the candidates, it computes two proxy losses for the unobservable true CATE estimation error, namely the R-loss (Nie and Wager 2021) and the DR-loss (Kennedy 2023; Mahajan et al. 2024), and also reports the factual loss (the prediction error for the observed outcomes). Because these steps are common to all candidates, differences in loss reflect differences in the candidates’ estimates.

Users then either select the model with the lowest value of a chosen loss using `cf_select` or combine the out-of-fold estimates of several models using `cf_ensemble`. The ensemble can be a simple average; a softmax-weighted average; stacking, that is, a weighted combination whose weights are learned to minimize the R-loss (Nie and Wager 2021; Han and Wu 2022) or the DR-loss (Han and Wu 2022); or an average of the models that agree most closely (Machluf et al. 2024).

### 2.3 Evaluation, interpretation and downstream analysis

After model selection or ensembling, the following functions evaluate and interpret the CATE estimates and support downstream analysis (Fig. 1D). Individual effects are unobservable, but a per-patient pseudo-outcome that equals the true CATE on average can be constructed from the observed outcomes; `cf_evaluate` uses this pseudo-outcome to assess the estimates. The rank-weighted average treatment effect (RATE; Yadlowsky et al. 2025) and sorted group average treatment effects (GATES; Chernozhukov et al. 2025) average the pseudo-outcome within groups of patients ranked by the estimates to test whether patients with larger estimates have larger true effects. The best linear predictor (BLP; Chernozhukov et al. 2025) regresses the pseudo-outcome on the estimates to quantify the linear relationship between the true effects and the estimates; a slope of 1 indicates correctly scaled estimates, and a slope of 0 indicates no such relationship. To identify which molecular features are associated with variation in the estimated CATE, and in which direction, `cf_interpret` uses SHAP values (Lundberg and Lee 2017) of a surrogate model that predicts the estimated CATE from the features. `cf_rule` constructs a treatment rule and estimates its policy value, that is, the mean outcome if patients were treated according to the rule (Athey and Wager 2021), and compares it with treating all or no patients. `cf_benefit_groups` assigns patients to benefit groups on the basis of their out-of-fold CATE estimates, and `cf_attach_results` writes the estimates and groups back to the original `SummarizedExperiment` for use in established Bioconductor workflows such as differential expression and gene-set analysis.

## 3 Application

To demonstrate the functionality of `CATEomics`, we analyzed gene expression data from baseline lesional skin biopsies of 98 patients with psoriasis (74 treated with brodalumab and 24 with placebo; GSE117468; Tomalin et al. 2020). The outcome was the negated week-12 Psoriasis Area and Severity Index (PASI; a severity score in which higher values indicate more severe disease), so that a positive effect indicates a lower PASI under brodalumab. The omics features were the expression data, and the clinical covariates were baseline PASI, age and body mass index. The analysis settings and code are given in Listing 1. In the scoreboard returned by `cf_compare`, the X-learner had the lowest R-loss, 7.6% lower than that obtained by predicting the same effect for all patients (Table 1a). R-loss-based stacking with `cf_ensemble` placed most of its weight on the X- and DR-learners (Supplementary Table S1).

**Table 1.**
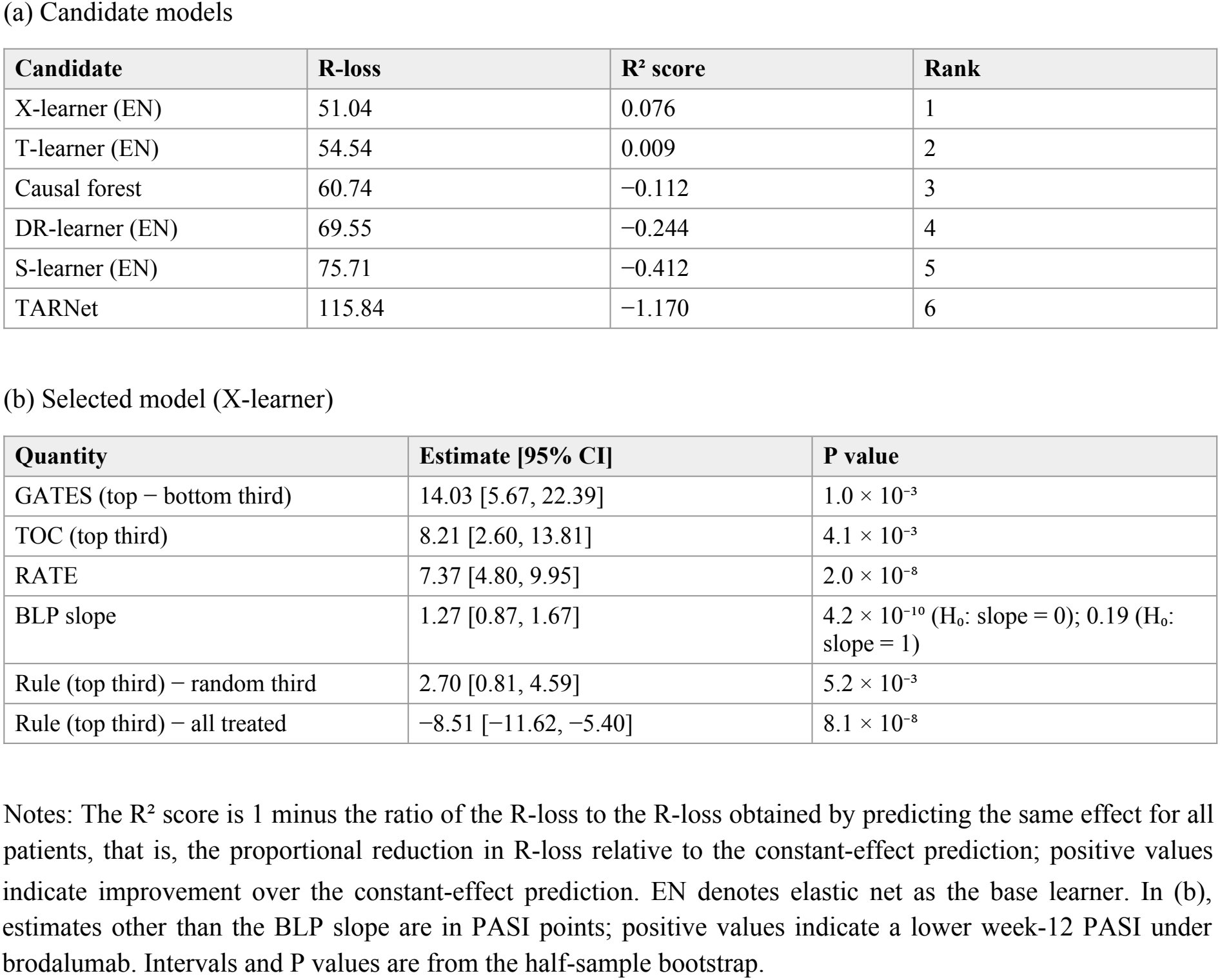
Comparison of candidate models and evaluation of the selected model.

| **(a) Candidate models** |  |  |  |
|---|---|---|---|
| **Candidate** | **R-loss** | **R² score** | **Rank** |
| X-learner (EN) | 51.04 | 0.076 | 1 |
| T-learner (EN) | 54.54 | 0.009 | 2 |
| Causal forest | 60.74 | −0.112 | 3 |
| DR-learner (EN) | 69.55 | −0.244 | 4 |
| S-learner (EN) | 75.71 | −0.412 | 5 |
| TARNet | 115.84 | −1.170 | 6 |
| (b) Selected model (X-learner) |  |  |  |
| **Quantity** | **Estimate [95% CI]** | **P value** |  |
| GATES (top − bottom third) | 14.03 [5.67, 22.39] | 1.0 × 10<sup>−3</sup> |  |
| TOC (top third) | 8.21 [2.60, 13.81] | 4.1 × 10<sup>−3</sup> |  |
| RATE | 7.37 [4.80, 9.95] | 2.0 × 10<sup>−8</sup> |  |
| BLP slope | 1.27 [0.87, 1.67] | 4.2 × 10<sup>−10</sup> (H<sub>0</sub>: slope = 0); 0.19 (H<sub>0</sub>: slope = 1) |  |
| Rule (top third) − random third | 2.70 [0.81, 4.59] | 5.2 × 10<sup>−3</sup> |  |
| Rule (top third) − all treated | −8.51 [−11.62, −5.40] | 8.1 × 10<sup>−8</sup> |  |

Notes: The R² score is 1 minus the ratio of the R-loss to the R-loss obtained by predicting the same effect for all patients, that is, the proportional reduction in R-loss relative to the constant-effect prediction; positive values indicate improvement over the constant-effect prediction. EN denotes elastic net as the base learner. In (b), estimates other than the BLP slope are in PASI points; positive values indicate a lower week-12 PASI under brodalumab. Intervals and P values are from the half-sample bootstrap.

The following downstream results were obtained with the X-learner selected by `cf_select`. Using `cf_evaluate` and the pseudo-outcome constructed from the observed outcomes, we assessed how well the estimates agreed with the true effects (Table 1b; Supplementary Table S2 and Fig. S1). When patients were divided into thirds by their estimates, the average effect in the third with the largest estimates, that is, the patients predicted to benefit most, was 14.0 PASI points larger than that in the third with the smallest estimates (GATES). The average treatment effect in the top third exceeded the overall average treatment effect by 8.2 PASI points, corresponding to the targeting operator characteristic (TOC) evaluated at the top-third cutoff. The RATE, a weighted integral of the TOC curve across different proportions of patients prioritized for treatment, was 7.4 PASI points. These results suggest that patients with larger estimates also had larger true effects. The BLP slope of 1.27 was significantly greater than 0, indicating that the estimates were related to the true effects, and did not differ significantly from 1, indicating that their scale was consistent with that of the true effects.

The rule from `cf_rule`, which treats the third of patients with the largest estimated effects, outperformed treating a random third of patients but did not outperform treating all patients (Table 1b; Supplementary Table S3). In the XGBoost surrogate used by `cf_interpret`, the COCH probe, baseline PASI and the ELOVL3 probe had the largest mean absolute SHAP values; higher feature values tended to contribute to larger estimated CATEs (Supplementary Table S4). `cf_benefit_groups` defined tertile benefit groups, which `cf_attach_results` added to the original object. A `limma` analysis (Ritchie et al. 2015) based on these groups detected 668 probes differentially expressed between the highest and lowest benefit groups (FDR < 0.05; Supplementary Table S5). In gene set variation analysis (GSVA; Hänzelmann et al. 2013), the scores of the Hallmark gene sets HEME_METABOLISM, PI3K_AKT_MTOR_SIGNALING and INTERFERON_ALPHA_RESPONSE (Liberzon et al. 2015) were negatively correlated with the CATE estimates (FDR < 0.05; Supplementary Table S6). These molecular associations are descriptive because the CATE estimates and benefit groups were derived from the same expression data.

**Listing 1.**
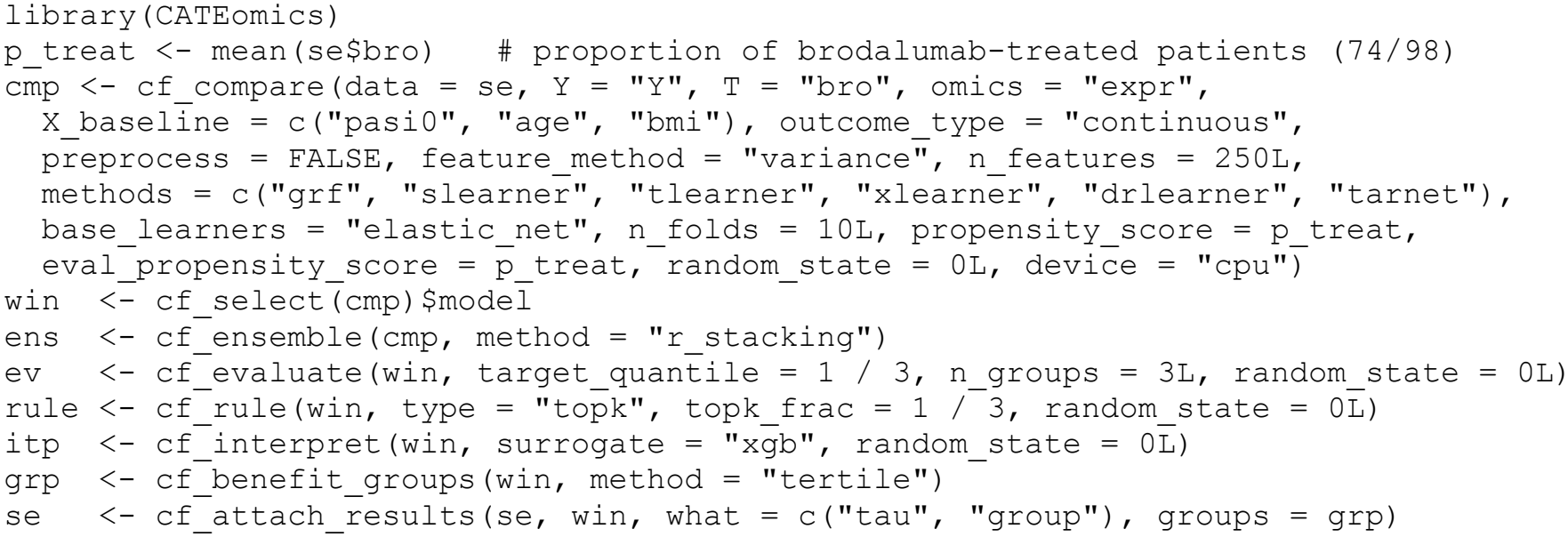
Code for the analysis in Section 3. The object `se` is a `SummarizedExperiment` built from GSE117468, with the expression matrix in the assay `expr` and the outcome, treatment and clinical covariates in `colData`.

## 4 Conclusion

`CATEomics` provides comparison, selection and ensembling of CATE estimators as a single reproducible workflow for omics data. Future development aims to add support for time-to-event outcomes and to extend support for multi-omics integration with `MultiAssayExperiment`.

## Author contributions

Takeru Furuta: Software, Data curation, Formal analysis, Validation, Visualization, Writing – original draft. Koichi Himori: Methodology, Validation, Writing – review & editing. Bingyuan Zhang: Methodology, Validation, Writing – review & editing. Akihiro Kitani: Methodology, Validation, Writing – review & editing. Yusuke Matsui: Conceptualization, Software, Writing – review & editing, Supervision, Funding acquisition.

## Data availability

The gene expression and clinical data analysed in this study are available from the Gene Expression Omnibus under accession GSE117468. The GSE117468 analysis scripts and results (v1.0.0) are archived on Zenodo at https://doi.org/10.5281/zenodo.24189811.

