## Supplementary Material for "CATEomics: model selection and ensembling for heterogeneous treatment effect estimation from high-dimensional omics data"

### Overview

This Supplementary Material describes the analysis in Section 3 of the main text. The package vignette describes its usage. The GSE117468 analysis scripts and results (v1.0.0) are archived on Zenodo at https://doi.org/10.5281/zenodo.24189811.

### 1 Main functions

| Function | Description |
|---|---|
| `cf_compare` | Fits the candidate models with common preprocessing and data partitions, and scores their out-of-fold estimates with the R-loss, DR-loss and factual loss |
| `cf_select` | Selects the candidate with the lowest value of a chosen loss |
| `cf_ensemble` | Combines the out-of-fold estimates of the candidates |
| `cf_fit` | Fits a single CATE estimator |
| `predict` | Returns the estimates from a fitted model and applies the model to new patients |
| `cf_evaluate` | Evaluates the estimates against the pseudo-outcome (TOC, RATE, GATES and BLP) |
| `cf_rule` | Constructs a treatment rule from the estimates and estimates its policy value |
| `cf_interpret` | Interprets the estimates with SHAP values of a surrogate model |
| `cf_benefit_groups` | Assigns patients to benefit groups on the basis of their out-of-fold estimates |
| `cf_attach_results` | Writes the estimates and benefit groups back to the original `SummarizedExperiment` |
| `cf_plot_*` | Plot the above results |

### 2 Data

We used GSE117468 from the Gene Expression Omnibus (GEO; Tomalin et al. 2020), the biopsy substudy of the AMAGINE-2 and AMAGINE-3 trials (platform GPL570). We analyzed baseline lesional skin biopsies from patients treated with brodalumab (140 mg or 210 mg every 2 weeks; the two doses were pooled) or placebo who had complete data on week-12 PASI, baseline PASI, age and body mass index: 98 patients (74 brodalumab and 24 placebo).

| Step | n |
|---|---|
| All samples in the series | 565 |
| Baseline | 256 |
| + Lesional skin | 128 |
| + Brodalumab (either dose) or placebo | 112 |
| + Complete week-12 PASI, baseline PASI, age and body mass index | 98 |

The 14 patients without week-12 PASI (12 brodalumab and 2 placebo) were excluded. The outcome was Y = −(week-12 PASI), so that larger values are better. The clinical covariates were baseline PASI, age and body mass index. The expression data (53,951 probes) were used as provided by GEO (log2 scale).

#### 3 Analysis settings

**Preprocessing.** Because the expression data had already been preprocessed (log2 scale), no normalization was applied within the package. The 250 probes with the largest variance were selected, and the 253 features, including the clinical covariates, were centered and scaled.

**Data partitions and propensity score.** A single ten-fold partition, stratified by treatment, was shared by all candidate models (`random_state` = 0). The propensity score was set to the known constant 74/98.

**Losses.** The candidate models were scored with the R-loss, DR-loss and factual loss. The nuisance models for these losses were fitted separately from the candidates with a super learner (van der Laan et al. 2007) whose library comprised the sample mean, ridge regression, elastic net, lasso, random forest and gradient boosting (depths 2 and 4). The $R^2$ score is 1 minus the ratio of a loss to its baseline loss, defined as the minimum of the same loss without covariates. It is thus the proportional reduction in loss relative to prediction without covariates: the constant-effect prediction for the R-loss and DR-loss, and the mean of each treatment arm for the factual loss. Positive values indicate improvement over this prediction.

**Candidate models.** The candidates were the causal forest, the S-, T-, X- and DR-learners, and TARNet. For the S-, T-, X- and DR-learners, all regression models, including the second stage of the X-learner and the final stage of the DR-learner, were elastic nets (`ElasticNetCV` in `scikit-learn`, with an L1 ratio of 0.5 and the penalty strength chosen by five-fold cross-validation). All other hyperparameters were the package defaults.

**Selection and ensembling.** The candidate with the lowest R-loss was selected. In R-loss-based stacking, non-negative weights, not constrained to sum to one, were estimated without an intercept (Nie and Wager 2021; Han and Wu 2022).

**Evaluation.** The pseudo-outcome was the doubly robust score constructed from the same nuisance models and propensity score as the losses. TOC was evaluated for the third of patients with the largest estimates, and GATES for tertiles of the estimates. RATE was the AUTOC, computed as the mean TOC over treated fractions $q = 0.1, 0.2, \ldots, 1.0$, and BLP was the slope of the regression of the pseudo-outcome on the estimates. Intervals and two-sided P values were obtained from a half-sample bootstrap (200 replicates) with a normal approximation.

**Treatment rule.** The rule treated the third of patients with the largest estimates. The policy-value curve was computed for treated fractions from 0.1 to 1.0.

**Interpretation.** Ridge regression and XGBoost surrogate models were fitted, and the XGBoost surrogate, which had the higher held-out $R^2$ in five-fold cross-validation, was reported.

**Benefit groups.** Patients were assigned to tertiles of their out-of-fold estimates.

**Gene annotation.** Probes were mapped to gene symbols with `hgu133plus2.db` (the first symbol was used when a probe mapped to several). For GSVA, each symbol was represented by its probe with the largest variance.

**Downstream analysis.** The benefit groups and the estimates written back by `cf_attach_results` were used directly in established Bioconductor workflows. The highest and lowest benefit groups (33 patients each) were compared with `limma`, adjusting for baseline PASI, age and body mass index, with Benjamini–Hochberg (BH) adjustment. For GSVA, the Spearman correlation between the score of each of the 50 Hallmark gene sets (Liberzon et al. 2015) and the estimates was computed, with BH adjustment. Because the benefit groups and the estimates were derived from the same expression data, these results are descriptive.

### 4 Computing environment

The analysis was run with R 4.6.0 (R Core Team 2026), Bioconductor 3.23 (Huber et al. 2015), `CATEomics` 0.99.0 and Python 3.12.3 (set up automatically through `basilisk`) on Ubuntu 24.04 with 28 CPU cores. The main packages are listed below; all versions are recorded in `session_info.txt` in the Zenodo deposit.

| Purpose | Package (version) |
|---|---|
| CATE estimation | `grf` 2.6.1 (Athey et al. 2019); `EconML` 0.16.0 (Battocchi et al. 2019); `CATENets` 0.2.4 (Curth and van der Schaar 2021); `scikit-learn` 1.6.1 (Pedregosa et al. 2011); `PyTorch` 2.12.0 (Paszke et al. 2019) |
| Learners and nuisance models | `glmnet` 5.0 (Friedman et al. 2010); `ranger` 0.18.0 (Wright and Ziegler 2017); `xgboost` 3.2.1.1 (R) and 2.1.4 (Python) (Chen and Guestrin 2016) |
| Interpretation | `shap` 0.48.0 (Lundberg and Lee 2017); `shapviz` 0.10.3 |
| Python environment | `basilisk` 1.24.0 (Lun 2022) |
| Data | `SummarizedExperiment` 1.42.0; `hgu133plus2.db` 3.13.0; `AnnotationDbi` 1.74.0 |
| Downstream analysis | `limma` 3.68.3 (Ritchie et al. 2015); `GSVA` 2.6.2 (Hänzelmann et al. 2013); `msigdbr` 26.1.0 |
| Plotting | `ggplot2` 4.0.3 (Wickham 2016); `patchwork` 1.3.2 |

### Supplementary tables

**Table S1.** Losses and $R^2$ scores of the six candidate models, and the weights of R-loss-based stacking.

| Candidate | R-loss | $R^2$ score | DR-loss | DR $R^2$ score | Factual loss | Factual $R^2$ score | Stacking weight |
|---|---|---|---|---|---|---|---|
| X-learner (EN) | 51.04 | 0.076 | 314.50 | 0.164 | 59.05 | 0.318 | 0.618 |
| T-learner (EN) | 54.54 | 0.009 | 330.19 | 0.118 | 59.05 | 0.318 | 0.000 |
| Causal forest | 60.74 | −0.112 | 394.45 | −0.046 | 87.47 | −0.006 | 0.000 |
| DR-learner (EN) | 69.55 | −0.244 | 288.96 | 0.222 | 96.75 | −0.145 | 0.166 |
| S-learner (EN) | 75.71 | −0.412 | 558.86 | −0.470 | 80.77 | 0.034 | 0.000 |
| TARNet | 115.84 | −1.170 | 737.61 | −0.945 | 177.46 | −1.098 | 0.000 |
| Sum of weights | | | | | | | 0.784 |

Notes: EN denotes elastic net as the base learner. The X- and T-learners have the same factual loss because the first stage of the X-learner consists of the same arm-wise outcome models as the T-learner. The stacking weights are not constrained to sum to one.

**Table S2.** Evaluation of the selected model and the ensemble.

| Model | Quantity | Estimate [95% CI] | SE | P value | Null hypothesis |
|---|---|---|---|---|---|
| X-learner (EN) | TOC (top third) | 8.21 [2.60, 13.81] | 2.86 | $4.1 \times 10^{-3}$ | 0 |
| X-learner (EN) | RATE (AUTOC) | 7.37 [4.80, 9.95] | 1.31 | $2.0 \times 10^{-8}$ | 0 |
| X-learner (EN) | BLP slope | 1.27 [0.87, 1.67] | 0.20 | $4.2 \times 10^{-10}$ | slope = 0 |
| X-learner (EN) | BLP slope | 1.27 [0.87, 1.67] | 0.20 | 0.19 | slope = 1 |
| X-learner (EN) | GATES (top − bottom third) | 14.03 [5.67, 22.39] | 4.27 | $1.0 \times 10^{-3}$ | 0 |
| X-learner (EN) | GATES group 1 (bottom third) | 10.79 [7.95, 13.64] | — | — | — |
| X-learner (EN) | GATES group 2 (middle third) | 14.24 [4.99, 23.49] | — | — | — |
| X-learner (EN) | GATES group 3 (top third) | 24.82 [16.84, 32.80] | — | — | — |
| Ensemble | TOC (top third) | 13.20 [7.18, 19.21] | 3.07 | $1.7 \times 10^{-5}$ | 0 |
| Ensemble | RATE (AUTOC) | 7.87 [4.96, 10.79] | 1.49 | $1.2 \times 10^{-7}$ | 0 |
| Ensemble | BLP slope | 1.59 [1.11, 2.07] | 0.24 | $8.5 \times 10^{-11}$ | slope = 0 |
| Ensemble | BLP slope | 1.59 [1.11, 2.07] | 0.24 | 0.016 | slope = 1 |
| Ensemble | GATES (top − bottom third) | 19.52 [10.03, 29.01] | 4.84 | $5.5 \times 10^{-5}$ | 0 |
| Ensemble | GATES group 1 (bottom third) | 10.03 [6.71, 13.35] | — | — | — |
| Ensemble | GATES group 2 (middle third) | 10.15 [2.44, 17.86] | — | — | — |
| Ensemble | GATES group 3 (top third) | 29.55 [20.60, 38.50] | — | — | — |

Notes: Estimates other than the BLP slope are in PASI points; positive values indicate a lower week-12 PASI under brodalumab. GATES group values are mean pseudo-outcomes. The ensemble is R-loss-based stacking. Intervals and two-sided P values are from the half-sample bootstrap.

**Table S3.** Policy values of the treatment rule constructed from the selected model.

(a) Policy values and contrasts

| Policy or contrast | Estimate [95% CI] | SE | P value |
|---|---|---|---|
| Rule (top third; 32 patients treated) | −11.87 [−14.77, −8.97] | 1.48 | — |
| Random third | −14.57 [−17.13, −12.00] | 1.31 | — |
| All treated* | −3.36 [−5.07, −1.65] | 0.87 | — |
| None treated* | −20.00 [−23.73, −16.27] | 1.90 | — |
| Rule − random third | 2.70 [0.81, 4.59] | 0.97 | $5.2 \times 10^{-3}$ |
| Rule − all treated | −8.51 [−11.62, −5.40] | 1.59 | $8.1 \times 10^{-8}$ |

| Policy or contrast | Estimate [95% CI] | SE | P value |
|---|---|---|---|
| Rule − none treated | 8.13 [5.53, 10.74] | 1.33 | $9.4 \times 10^{-10}$ |

(b) Policy-value curve

| Treated fraction | Patients treated | Policy value [95% CI] |
|---|---|---|
| 0.1 | 9 | −16.35 [−20.11, −12.60] |
| 0.2 | 19 | −14.38 [−17.60, −11.15] |
| 0.3 | 29 | −11.87 [−14.93, −8.81] |
| 0.4 | 39 | −10.30 [−13.26, −7.35] |
| 0.5 | 49 | −7.53 [−9.93, −5.14] |
| 0.6 | 58 | −7.27 [−9.32, −5.22] |
| 0.7 | 68 | −6.61 [−8.43, −4.79] |
| 0.8 | 78 | −5.38 [−7.22, −3.53] |
| 0.9 | 88 | −3.96 [−5.70, −2.22] |
| 1.0 | 98 | −3.36 [−5.06, −1.66] |

Notes: The policy value is the estimated mean of Y (the negated week-12 PASI) if patients were treated according to the policy; larger values are better. Intervals and P values are from the half-sample bootstrap, except for the policies marked *, which are doubly robust arm means with influence-function intervals.

**Table S4.** Interpretation with the surrogate model.

(a) Top 20 features of the XGBoost surrogate

| Rank | Feature | Gene symbol | Mean absolute SHAP value | Direction |
|---|---|---|---|---|
| 1 | 205229_s_at | COCH | 1.262 | + |
| 2 | Baseline PASI | — | 0.824 | + |
| 3 | 234513_at | ELOVL3 | 0.705 | + |
| 4 | 1560756_at | — | 0.615 | − |
| 5 | 210797_s_at | OASL | 0.521 | − |
| 6 | 1552620_at | SPRR4 | 0.497 | + |
| 7 | 206367_at | REN | 0.469 | − |
| 8 | 234700_s_at | RNASE7 | 0.359 | − |
| 9 | 221872_at | RARRES1 | 0.292 | + |
| 10 | 237120_at | KRT77 | 0.272 | + |
| 11 | 216935_at | LINC00302 | 0.250 | − |
| 12 | 221404_at | IL36A | 0.237 | − |
| 13 | 219270_at | CHAC1 | 0.237 | − |
| 14 | 231583_at | KRT74 | 0.162 | − |
| 15 | 239430_at | IGFL1 | 0.156 | − |
| 16 | 1561754_at | — | 0.141 | + |
| 17 | 237732_at | PRR9 | 0.130 | − |
| 18 | 223720_at | SPINK7 | 0.125 | − |
| 19 | 231033_at | DSC2 | 0.124 | − |
| 20 | 207908_at | KRT2 | 0.116 | − |

(b) Fidelity (held-out $R^2$, five-fold)

| Surrogate | Held-out $R^2$ |
|---|---|
| Ridge regression | 0.636 |
| XGBoost (reported) | 0.664 |

Notes: Direction is the sign of the correlation between a feature and its SHAP values; + indicates that higher values of the feature tend to contribute to larger estimated CATEs. — denotes probes without a gene symbol.

**Table S5.** `limma` analysis of the highest versus lowest benefit group.

(a) Top 50 probes

| Rank | Probe | Gene symbol | logFC (high − low) | Average expression | P value | Adjusted P value |
|---|---|---|---|---|---|---|
| 1 | 205029_s_at | FABP7 | 2.12 | 6.61 | $2.3 \times 10^{-9}$ | $5.2 \times 10^{-5}$ |
| 2 | 205030_at | FABP7 | 2.81 | 6.42 | $2.9 \times 10^{-9}$ | $5.2 \times 10^{-5}$ |
| 3 | 211113_s_at | ABCG1 | −0.84 | 6.20 | $2.9 \times 10^{-9}$ | $5.2 \times 10^{-5}$ |
| 4 | 204567_s_at | ABCG1 | −0.82 | 10.18 | $5.8 \times 10^{-8}$ | $7.3 \times 10^{-4}$ |
| 5 | 217013_at | AZGP1P1 | −1.08 | 5.64 | $6.8 \times 10^{-8}$ | $7.3 \times 10^{-4}$ |
| 6 | 228568_at | MYZAP | −0.90 | 8.64 | $9.8 \times 10^{-8}$ | $8.8 \times 10^{-4}$ |
| 7 | 203703_s_at | TTLL4 | 0.62 | 4.33 | $2.7 \times 10^{-7}$ | $2.0 \times 10^{-3}$ |
| 8 | 211651_s_at | LAMB1 | 0.43 | 8.40 | $3.2 \times 10^{-7}$ | $2.1 \times 10^{-3}$ |
| 9 | 207213_s_at | USP2 | −0.95 | 5.16 | $4.3 \times 10^{-7}$ | $2.6 \times 10^{-3}$ |
| 10 | 231663_s_at | ARG1 | −1.61 | 8.21 | $5.7 \times 10^{-7}$ | $2.6 \times 10^{-3}$ |
| 11 | 1553857_at | IGSF22 | −0.59 | 7.22 | $5.8 \times 10^{-7}$ | $2.6 \times 10^{-3}$ |
| 12 | 1557609_s_at | TBC1D12 | −0.92 | 5.72 | $5.8 \times 10^{-7}$ | $2.6 \times 10^{-3}$ |
| 13 | 212884_x_at | APOE | 0.65 | 6.64 | $6.5 \times 10^{-7}$ | $2.6 \times 10^{-3}$ |
| 14 | 209583_s_at | CD200 | 0.76 | 5.22 | $6.8 \times 10^{-7}$ | $2.6 \times 10^{-3}$ |
| 15 | 230527_at | LOC101926907 | 0.52 | 5.57 | $7.5 \times 10^{-7}$ | $2.6 \times 10^{-3}$ |
| 16 | 204416_x_at | APOC1 | 1.38 | 7.40 | $8.1 \times 10^{-7}$ | $2.6 \times 10^{-3}$ |
| 17 | 234664_at | LINC01347 | −0.30 | 5.50 | $8.7 \times 10^{-7}$ | $2.6 \times 10^{-3}$ |
| 18 | 229740_at | SMIM5 | −0.68 | 9.28 | $8.7 \times 10^{-7}$ | $2.6 \times 10^{-3}$ |
| 19 | 215150_at | YOD1 | −1.51 | 6.91 | $1.0 \times 10^{-6}$ | $2.8 \times 10^{-3}$ |
| 20 | 223298_s_at | NT5C3A | −0.92 | 10.35 | $1.2 \times 10^{-6}$ | $3.1 \times 10^{-3}$ |
| 21 | 237296_at | — | −1.05 | 4.88 | $1.2 \times 10^{-6}$ | $3.1 \times 10^{-3}$ |
| 22 | 206113_s_at | RAB5A | −0.81 | 7.67 | $1.2 \times 10^{-6}$ | $3.1 \times 10^{-3}$ |
| 23 | 213553_x_at | APOC1 | 1.15 | 7.41 | $1.3 \times 10^{-6}$ | $3.1 \times 10^{-3}$ |
| 24 | 221416_at | PLA2G2F | −0.92 | 4.78 | $1.4 \times 10^{-6}$ | $3.1 \times 10^{-3}$ |
| 25 | 1567100_at | — | 0.95 | 4.55 | $1.4 \times 10^{-6}$ | $3.1 \times 10^{-3}$ |
| 26 | 1565525_a_at | TCP11L2 | −0.86 | 5.41 | $1.7 \times 10^{-6}$ | $3.3 \times 10^{-3}$ |
| 27 | 224806_at | TRIM25 | −0.57 | 10.42 | $1.7 \times 10^{-6}$ | $3.3 \times 10^{-3}$ |
| 28 | 242103_at | TMEM86A | −1.01 | 6.63 | $1.7 \times 10^{-6}$ | $3.3 \times 10^{-3}$ |
| 29 | 239770_at | ESYT3 | −1.05 | 7.02 | $1.8 \times 10^{-6}$ | $3.4 \times 10^{-3}$ |
| 30 | 216701_at | KPLCE | −1.67 | 9.23 | $1.9 \times 10^{-6}$ | $3.5 \times 10^{-3}$ |
| 31 | 227234_at | IPO5P1 | 0.45 | 5.54 | $2.1 \times 10^{-6}$ | $3.5 \times 10^{-3}$ |
| 32 | 216718_at | LINC00302 | −1.63 | 8.41 | $2.1 \times 10^{-6}$ | $3.5 \times 10^{-3}$ |
| 33 | 206656_s_at | APMAP | 0.58 | 9.48 | $2.1 \times 10^{-6}$ | $3.5 \times 10^{-3}$ |
| 34 | 203770_s_at | STS | −1.27 | 7.02 | $2.2 \times 10^{-6}$ | $3.5 \times 10^{-3}$ |
| 35 | 224051_at | PFKFB2 | −1.13 | 5.36 | $2.3 \times 10^{-6}$ | $3.5 \times 10^{-3}$ |
| 36 | 238455_at | PLXDC2 | −0.45 | 7.47 | $2.4 \times 10^{-6}$ | $3.5 \times 10^{-3}$ |
| 37 | 208023_at | TNFRSF4 | −0.65 | 7.23 | $2.4 \times 10^{-6}$ | $3.5 \times 10^{-3}$ |
| 38 | 238213_at | MIR5188 | −0.78 | 3.64 | $2.5 \times 10^{-6}$ | $3.5 \times 10^{-3}$ |
| 39 | 235857_at | KCTD11 | −0.64 | 9.15 | $2.6 \times 10^{-6}$ | $3.6 \times 10^{-3}$ |
| 40 | 205229_s_at | COCH | 2.01 | 5.90 | $2.6 \times 10^{-6}$ | $3.6 \times 10^{-3}$ |
| 41 | 236069_at | — | −0.96 | 9.38 | $3.1 \times 10^{-6}$ | $4.0 \times 10^{-3}$ |
| 42 | 209089_at | RAB5A | −0.35 | 11.75 | $3.4 \times 10^{-6}$ | $4.3 \times 10^{-3}$ |
| 43 | 223123_s_at | PITHD1 | −0.51 | 3.86 | $3.5 \times 10^{-6}$ | $4.3 \times 10^{-3}$ |
| 44 | 220431_at | TMPRSS11E | 1.34 | 4.52 | $3.5 \times 10^{-6}$ | $4.3 \times 10^{-3}$ |
| 45 | 227309_at | YOD1 | −0.79 | 11.21 | $4.0 \times 10^{-6}$ | $4.7 \times 10^{-3}$ |
| 46 | 1555383_a_at | POF1B | −0.97 | 8.11 | $4.0 \times 10^{-6}$ | $4.7 \times 10^{-3}$ |
| 47 | 213260_at | FOXC1 | 0.82 | 9.09 | $4.2 \times 10^{-6}$ | $4.7 \times 10^{-3}$ |
| 48 | 228309_at | — | 0.57 | 9.92 | $4.2 \times 10^{-6}$ | $4.7 \times 10^{-3}$ |
| 49 | 225018_at | SPIRE1 | −0.66 | 7.82 | $4.3 \times 10^{-6}$ | $4.7 \times 10^{-3}$ |

| Rank | Probe | Gene symbol | logFC (high − low) | Average expression | P value | Adjusted P value |
|---|---|---|---|---|---|---|
| 50 | 217087_at | KPLCE | −1.63 | 10.28 | $4.3 \times 10^{-6}$ | $4.7 \times 10^{-3}$ |

(b) Summary

| Item | Number of probes |
|---|---|
| Tested | 53,951 |
| FDR < 0.05 | 668 |
| FDR < 0.10 | 1,602 |

Notes: The highest and lowest benefit groups are the top and bottom tertiles of the out-of-fold estimates (33 patients each). The model was adjusted for baseline PASI, age and body mass index. P values were adjusted by the Benjamini–Hochberg method.

**Table S6.** Spearman correlation between the GSVA scores of the Hallmark gene sets and the CATE estimates.

| Gene set | ρ | P value | FDR | Genes found / set size |
|---|---|---|---|---|
| HEME_METABOLISM | −0.403 | $4.6 \times 10^{-5}$ | $2.3 \times 10^{-3}$ | 199 / 200 |
| PI3K_AKT_MTOR_SIGNALING | −0.376 | $1.5 \times 10^{-4}$ | $3.8 \times 10^{-3}$ | 105 / 105 |
| INTERFERON_ALPHA_RESPONSE | −0.310 | $2.0 \times 10^{-3}$ | 0.033 | 97 / 97 |
| PROTEIN_SECRETION | −0.279 | $5.5 \times 10^{-3}$ | 0.060 | 96 / 96 |
| INTERFERON_GAMMA_RESPONSE | −0.277 | $6.0 \times 10^{-3}$ | 0.060 | 200 / 200 |
| TNFA_SIGNALING_VIA_NFKB | −0.251 | 0.013 | 0.11 | 200 / 200 |
| ESTROGEN_RESPONSE_LATE | −0.233 | 0.021 | 0.13 | 200 / 200 |
| UNFOLDED_PROTEIN_RESPONSE | −0.226 | 0.025 | 0.13 | 112 / 113 |
| UV_RESPONSE_UP | −0.223 | 0.028 | 0.13 | 158 / 158 |
| MTORC1_SIGNALING | −0.221 | 0.029 | 0.13 | 200 / 200 |
| P53_PATHWAY | −0.218 | 0.031 | 0.13 | 200 / 200 |
| IL6_JAK_STAT3_SIGNALING | −0.215 | 0.034 | 0.13 | 87 / 87 |
| APICAL_SURFACE | −0.214 | 0.035 | 0.13 | 44 / 44 |
| REACTIVE_OXYGEN_SPECIES_PATHWAY | −0.208 | 0.040 | 0.14 | 49 / 49 |
| MITOTIC_SPINDLE | −0.200 | 0.049 | 0.16 | 199 / 199 |
| COMPLEMENT | −0.192 | 0.059 | 0.18 | 200 / 200 |
| HYPOXIA | −0.190 | 0.061 | 0.18 | 198 / 200 |
| G2M_CHECKPOINT | −0.180 | 0.076 | 0.21 | 200 / 200 |
| GLYCOLYSIS | −0.175 | 0.084 | 0.22 | 200 / 200 |
| DNA_REPAIR | −0.169 | 0.097 | 0.24 | 150 / 150 |
| MYC_TARGETS_V1 | −0.154 | 0.13 | 0.29 | 200 / 200 |
| IL2_STAT5_SIGNALING | −0.151 | 0.14 | 0.29 | 199 / 199 |
| WNT_BETA_CATENIN_SIGNALING | −0.149 | 0.14 | 0.29 | 41 / 42 |
| EPITHELIAL_MESENCHYMAL_TRANSITION | 0.147 | 0.15 | 0.29 | 200 / 200 |
| ADIPOGENESIS | 0.147 | 0.15 | 0.29 | 200 / 200 |
| E2F_TARGETS | −0.145 | 0.15 | 0.29 | 199 / 200 |
| INFLAMMATORY_RESPONSE | −0.142 | 0.16 | 0.29 | 200 / 200 |
| NOTCH_SIGNALING | −0.141 | 0.17 | 0.29 | 32 / 32 |
| APOPTOSIS | −0.138 | 0.17 | 0.29 | 161 / 161 |
| MYOGENESIS | 0.138 | 0.18 | 0.29 | 200 / 200 |
| BILE_ACID_METABOLISM | 0.137 | 0.18 | 0.29 | 111 / 112 |
| XENOBIOTIC_METABOLISM | −0.133 | 0.19 | 0.30 | 199 / 200 |
| ALLOGRAFT_REJECTION | −0.132 | 0.20 | 0.30 | 200 / 200 |
| FATTY_ACID_METABOLISM | 0.130 | 0.20 | 0.30 | 156 / 158 |
| ANGIOGENESIS | 0.120 | 0.24 | 0.34 | 36 / 36 |

| Gene set | ρ | P value | FDR | Genes found / set size |
|---|---|---|---|---|
| KRAS_SIGNALING_UP | −0.111 | 0.28 | 0.39 | 200 / 200 |
| UV_RESPONSE_DN | 0.097 | 0.34 | 0.46 | 144 / 144 |
| TGF_BETA_SIGNALING | −0.088 | 0.39 | 0.51 | 54 / 54 |
| ANDROGEN_RESPONSE | −0.084 | 0.41 | 0.53 | 101 / 101 |
| KRAS_SIGNALING_DN | −0.078 | 0.44 | 0.56 | 199 / 200 |
| SPERMATOGENESIS | −0.068 | 0.50 | 0.62 | 134 / 135 |
| APICAL_JUNCTION | 0.057 | 0.57 | 0.68 | 200 / 200 |
| PEROXISOME | −0.055 | 0.59 | 0.69 | 104 / 104 |
| ESTROGEN_RESPONSE_EARLY | −0.050 | 0.63 | 0.71 | 200 / 200 |
| PANCREAS_BETA_CELLS | −0.045 | 0.66 | 0.74 | 40 / 40 |
| COAGULATION | −0.042 | 0.68 | 0.74 | 138 / 138 |
| MYC_TARGETS_V2 | 0.021 | 0.83 | 0.89 | 58 / 58 |
| OXIDATIVE_PHOSPHORYLATION | 0.009 | 0.93 | 0.97 | 198 / 200 |
| HEDGEHOG_SIGNALING | 0.004 | 0.97 | 0.98 | 36 / 36 |
| CHOLESTEROL_HOMEOSTASIS | −0.002 | 0.98 | 0.98 | 74 / 74 |

Notes: n = 98. FDR values are Benjamini–Hochberg-adjusted over the 50 gene sets. Genes found are those among the 21,202 gene symbols in the data.

### Supplementary figure

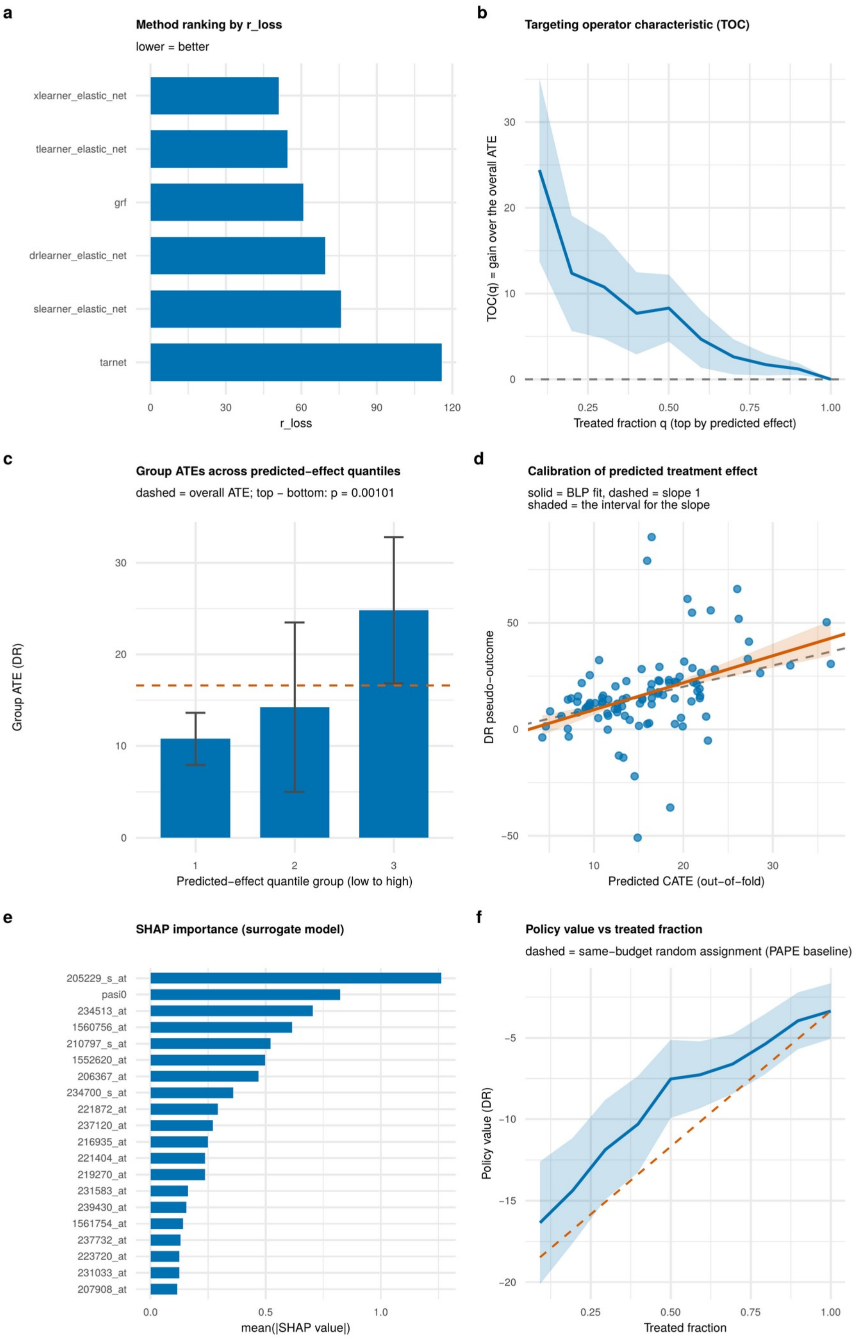


**Fig. S1.** Output of the `CATEomics` plotting functions for the analysis of GSE117468. (a) `cf_plot_compare`: R-loss of the six candidate models (lower is better). (b) `cf_plot_evaluate` (TOC): the amount by which the average treatment effect among the fraction q of patients with the largest estimates exceeds the overall average treatment effect; the band shows 95% confidence intervals. (c) `cf_plot_evaluate` (GATES): mean pseudo-outcome in each tertile of the estimates; error bars show 95% confidence intervals, and the dashed line shows the overall average treatment effect. (d) `cf_plot_evaluate` (BLP): pseudo-outcome of each patient against the out-of-fold estimate; the solid line shows the BLP fit (slope 1.27; 95% CI 0.87–1.67) and the shaded region the lines with slopes within this interval, and the dashed line shows a slope of 1 (perfect calibration). (e) `cf_plot_interpret`: top 20 features by mean absolute SHAP value of the XGBoost surrogate (gene symbols are given in Table S4). (f) `cf_plot_policy_curve`: policy value by treated fraction, treating patients with the largest estimates first; the band shows 95% confidence intervals, and the dashed line shows treatment of a random group of the same size.
